# Design, Structure, and Immunogenicity of a Soluble Prefusion-stabilized EBV gB Antigen

**DOI:** 10.1101/2025.05.19.654955

**Authors:** Ryan S. McCool, Cory M. Acreman, Abigail E. Powell, Sofia I. Picucci, Daniel J. Stieh, Jeremy Huynh, Hannah Caruso, Soyoon Park, Jessica O’Rear, Jui-Lin Chen, Brad A. Palanski, Patrick O. Byrne, Madeline R. Sponholtz, Chia-Wei Chou, Jeongryeol Kim, Julie E. Ledgerwood, Payton A.-B. Weidenbacher, Jason S. McLellan

## Abstract

Epstein-Barr virus (EBV), the causative agent of mononucleosis, is linked to over 140,000 annual cancer-related deaths globally and increases the risk of multiple sclerosis by up to 32-fold. As a herpesvirus, EBV establishes lifelong infection, and over 90% of U.S. adults are EBV-seropositive. Despite its significant disease burden, no approved EBV vaccines or therapeutics exist. Among EBV envelope glycoproteins, the fusion protein (gB) is strictly required for epithelial and B cell infection. Using a combination of AlphaFold-guided modeling, rational design, and ThermoMPNN-informed optimization, we engineered a stabilized prefusion gB variant, D2C3. This construct incorporates two inter-protomeric disulfide bonds and three cavity-filling substitutions, resulting in a melting temperature of 54 °C. Cryo-EM analysis of this construct allowed us to determine the prefusion structure of EBV gB, providing insights into the structural transitions required to adopt the postfusion conformation. Murine immunizations and depletion studies with human sera suggested a trend toward improved functional immunogenicity of D2C3 compared to postfusion gB. Collectively, these studies define engineering principles to stabilize class III fusion proteins, provide reagents to interrogate the human antibody response to EBV gB, and lay a foundation for further studies to develop EBV gB-based vaccine candidates.

## Introduction

Epstein-Barr virus (EBV), also known as *human γ-herpesvirus 4*, contributes to more than 140,000 cancer-related deaths annually^1^. EBV is the viral cause of mononucleosis, and recent studies have linked EBV seropositivity to a 26- to 32-fold increased risk of multiple sclerosis, highlighting its broad disease burden^2,3^. Like other members of *Herpesviridae*, EBV establishes lifelong infection, cycling between lytic stages—characterized by the production of infectious virions—and latent stages, where there is minimal viral transcription. Despite its significant clinical impact, no approved EBV vaccines or therapeutics exist.

Six glycoproteins on the viral envelope mediate EBV attachment and membrane fusion, with glycoprotein B (gB) acting as the essential fusion protein for infecting susceptible cell types, which include epithelial cells and B cells. EBV gB is translated as a single peptide decorated by eight N-linked and five O-linked glycans^4^. The gB monomer contains five intraprotomer disulfide bonds and trimerizes to form a metastable prefusion complex^5,6^. EBV gB is cleaved by furin, which enhances viral fusion^7^, although both the cleaved and uncleaved forms are present on mature virions^8^. Similar to other herpesvirus gB proteins, EBV gB comprises five structural domains (I–V)^9^. To facilitate membrane fusion and entry, gB transitions from a metastable prefusion conformation to a stable postfusion conformation^10,11^. This rearrangement is likely triggered by receptor engagement of the glycoprotein H/glycoprotein L (gH/gL) heterodimer complex for epithelial cell fusion and the gH/gL/glycoprotein 42 (gp42) heterotrimer complex for B cell fusion^12–14^.

Notably, membrane fusion can occur in the absence of other viral glycoproteins if gB is heat-treated or its cytoplasmic tail is truncated^14,15^. Moreover, antibodies targeting gB inhibit fusion across all susceptible cell types^16,17^.

Prefusion stabilization has enhanced immunogenicity of some class I viral fusion proteins, leading to substantially improved elicitation of neutralizing antibody titers^18–23^. This enhancement arises in part from increased production of prefusion-specific antibodies that block conformational rearrangements required for fusion. Unlike class I fusion proteins, which undergo major secondary structure rearrangements following triggering^19,24–28^, class III proteins such as gB primarily reorganize the spatial positioning of domains I–V with minimal changes to secondary structure^9–11^. Prior efforts to stabilize a soluble prefusion gB ectodomain from human cytomegalovirus (HCMV), a related β-herpesvirus, relied on engineered disulfide bonds^29^. However, this construct exhibited flexibility in domain I compared to a prefusion, membrane-anchored HCMV gB stabilized by crosslinkers and a fusion inhibitor^10,11^.

Additionally, soluble prefusion-like HCMV gB failed to elicit higher neutralizing antibody titers in immunized mice. Given that gB is highly conserved across *Herpesviridae*^30^, shares some structural features with class I fusion proteins^5^, and that gB is necessary and sufficient for EBV-mediated membrane fusion^14,15,17^, we aimed to stabilize EBV gB in its prefusion state and assess its immunogenicity as a vaccine antigen.

Here, we employed sequence alignments and AlphaFold2 structural predictions^31,32^ of prefusion EBV gB to guide stabilization efforts. We show that direct transfer of HCMV gB stabilizing substitutions proved insufficient, necessitating a broader design strategy. We then describe the engineering, structural validation, and immunogenic assessment of D2C3, a prefusion-stabilized EBV gB antigen optimized through computational modeling, rational design, and experimental screening^33,34^. Our findings demonstrate that D2C3 maintains a prefusion conformation, and we provide a high-resolution structure of the soluble prefusion gB ectodomain. Immunization of mice with D2C3 elicits comparable, if slightly improved, neutralizing antibody responses and preferentially depletes neutralizing antibodies from human sera obtained from EBV-seropositive individuals compared to postfusion gB. These results establish prefusion gB as a promising vaccine antigen and provide a framework for future studies on herpesvirus entry and immunogen design.

## Results

### Design and Initial Characterization of Single Substitution EBV gB Variants

The design of a soluble gB ‘base’ construct (EBV Base), in which prefusion-stabilizing substitutions would be assessed, was guided by previous work on a soluble postfusion EBV gB ectodomain construct^6^. Our base construct consisted of the EBV gB ectodomain (M81 strain, residues 1– 688) fused to a C-terminal T4 fibritin (foldon) trimerization motif, an HRV 3C recognition sequence, an octa-histidine tag, and a Twin-Strep tag (Fig. 1A; Supplementary Fig. 1A). To achieve sufficient yields, we substituted the EBV gB fusion loop residues (112-WY-113 and 193-WLIW-196) with the corresponding HSV-2 residues (177-HR-178 and 258-RVEA-261; Supplementary Table S1, Supplementary Figs. S1A, S2, S3). Consistent with previous findings, constructs lacking this substitution exhibited prohibitively low expression yields^5,35^. Additionally, replacing the furin cleavage site (428-RRRRR-432) between domains II and III with a GGSGG sequence improved homogeneity, as assessed by reducing SDS-PAGE (Supplementary Figs. S2, S3B). We also included the D220E substitution, present in the B95-8 strain of EBV, to maintain an EcoRI restriction site for molecular cloning (Supplementary Table S1, Figs. S2, S3). Using these modifications, we routinely observed yields of approximately 1.2 mg of EBV Base from 80 mL of transiently transfected FreeStyle 293 cell cultures after 6 days of culture.

**Figure 1.**
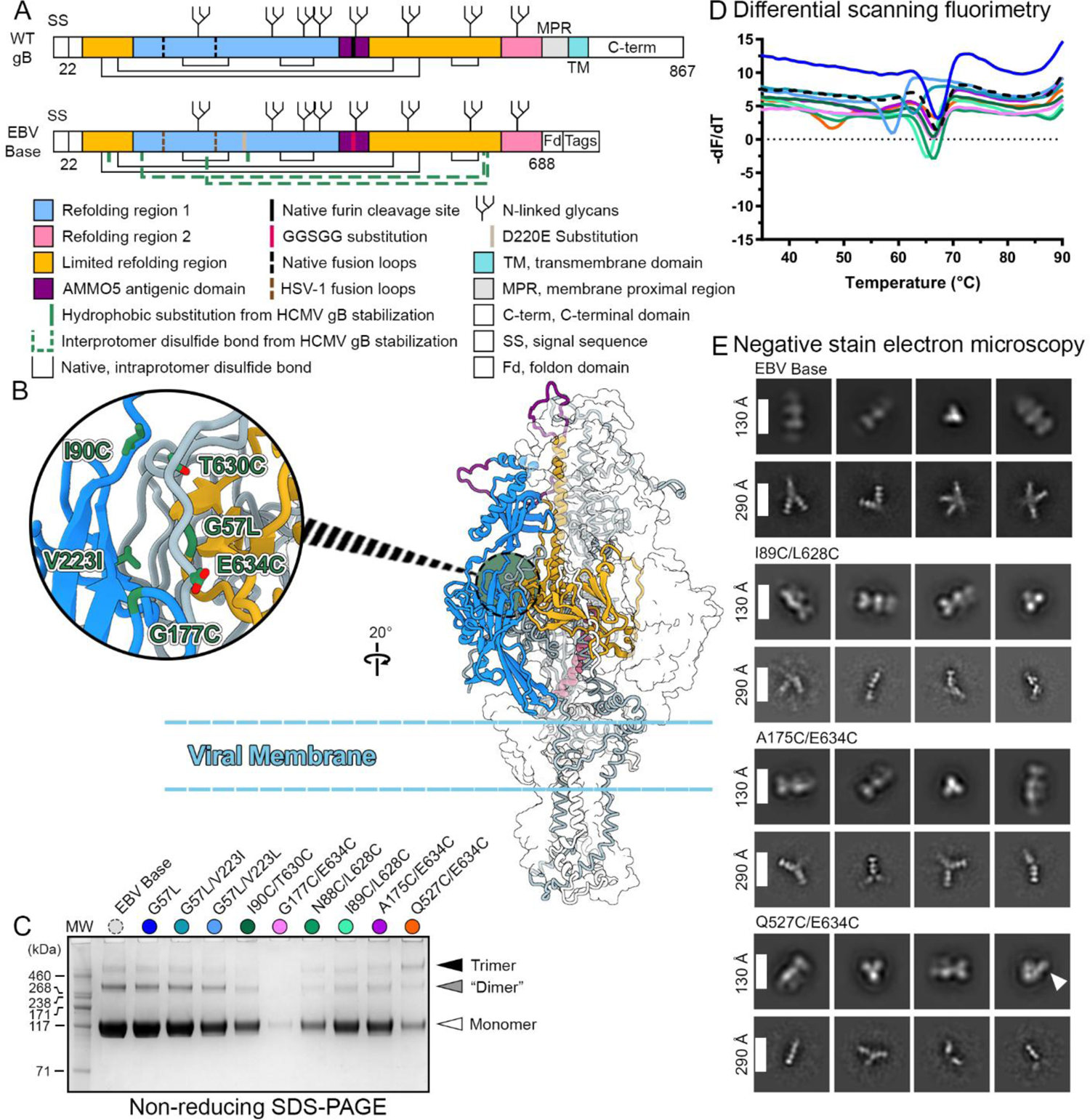
Characterization of EBV gB variants suggests subtle structural differences from HCMV gB. (A) Schematic of wildtype (WT) gB and ectodomain base construct (EBV Base). EBV Base contains modifications listed in the diagram. (B) AlphaFold2 model of prefusion WT gB. Trimeric prefusion HCMV gB (PDB ID: 7KDP) was used as a template. One protomer is colored as in (A) and shown as a ribbon, one is colored gray and shown as a trace of the α-carbon backbone, and one is shown as a transparent surface. Inset shows EBV gB residues homologous to those used to stabilize a prefusion-like HCMV gB ectodomain^29^. Residues are shown as green sticks with oxygen atoms shown in red. (C) Non-reducing SDS-PAGE analysis of gB variants. (D) DSF analysis of gB variant thermostability, colored as in (C). (E) nsEM 2D averages of top single-disulfide variants. White arrow highlights a splayed-out DI.

To stabilize gB in the prefusion conformation, we used sequence alignments (Supplementary Fig. S2) and an AlphaFold2 model––generated using the prefusion, membrane-anchored HCMV gB structure as a template––to inform our designs (PDB ID: 7KDP; Fig. 1B; Supplementary Figs. S1B, S2, S4)^11,31,36^. In HCMV gB, successful designs formed interprotomer disulfides that were detectable as monomer-to-trimer shifts on non-reducing SDS-PAGE. However, when applied to EBV gB, homologous designs from prefusion-like HCMV gB—I90C/T630C and G177C/E634C—failed to induce the expected monomer-to-trimer shift (Fig. 1C, Table 1). Similarly, the G57L/V223I variant, homologous to T100L/A267I in HCMV gB, failed to improve expression yields. These results suggested that subtle yet important structural differences exist between EBV and HCMV gB.

**Table 1.**
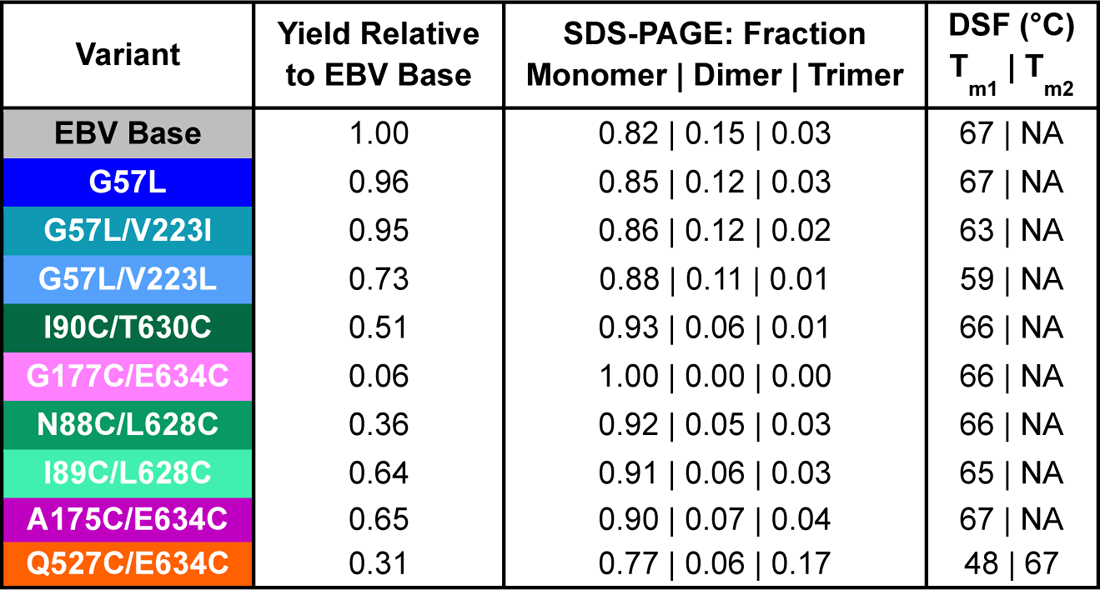
Characteristics of EBV gB single variants. All values were calculated from the non-reducing gel or the DSF profile in Fig. 1.

To further evaluate stabilization strategies, we employed a disulfide scanning approach to assess if residues adjacent to our initial target substitutions could more readily form stabilizing disulfide bonds. This identified several promising disulfides (N88C/L628C, I89C/L628C, A175C/E634C, and Q527C/E634C) capable of increasing the trimer-to-monomer band ratios observed on non-reducing SDS-PAGE (Fig. 1C; Table 1). Differential scanning fluorimetry (DSF) showed that EBV Base had a higher melting temperature (T_m_) than HCMV gB Base (67 vs. 59 °C, respectively; Fig. 1D)^29^. Most variants had T_m_ values within 3 °C of EBV Base, except G57L/V223I (63 °C) and G57L/V223L (59 °C). Notably, Q527C/E634C displayed a unique thermal profile, with a lower T_m_ (48 °C) and an additional thermal transition at 67 °C. Negative-stain electron microscopy (nsEM) revealed that the EBV Base construct as well as the single-disulfide variants exclusively adopted the postfusion conformation (Fig. 1E), displaying the characteristic crown (domain IV) and compact base (domain V) features of postfusion gB^6,37–39^. Collectively, these data demonstrated that multiple substitutions would be required to stabilize EBV gB in a prefusion conformation.

### Combinatorial Variant Designs Stabilize Prefusion gB

We next assessed combinatorial designs to evaluate whether pairing stabilizing disulfides could enhance trimer formation and prefusion stability. Although all variants incorporating multiple interprotomer disulfide bonds exhibited higher trimer content than their respective single variants, those containing N88C/L628C (C1 and C2) exhibited lower expression than those with I89C/L628C (C3 and C4) (Supplementary Table S2; Supplementary Fig. S5A). Additionally, double-disulfide variants containing Q527C/E634C (C2 and C4) improved the trimer-to-monomer ratio on non-reducing SDS-PAGE, whereas incorporating A175C/E634C (C1 and C3) enhanced thermostability of the first thermal transition (T _m1_) (Supplementary Fig. S5B). DSF analyses also revealed unique thermal transitions compared to EBV Base for all combinatorial designs except for C1.

As C2 and C4 showed no monomer bands on non-reducing SDS-PAGE, we evaluated these samples by nsEM. 2D class averages of both variants revealed protrusions from a globular core (Supplementary Fig. S5C, white arrows). These features are inconsistent with the expected compact prefusion conformation and are similar to those observed in the Q527C/E634C nsEM dataset (Fig. 1E, white arrow). Given that the disulfides in C2 and C4 were designed to covalently link domains II and IV to domain V, we hypothesized that these protrusions represent domain I, which may hinge outward at the ‘elbow’ connecting domain I to domain II (Supplementary Fig. S1B, black arrows). These results suggested that further stabilization of the variants containing the A175C/E634C disulfide bond (C1 and C3), which covalently links domain I to domain V, would be a more promising approach for prefusion-stabilization of gB.

We hypothesized that disulfide bonds incorporating L628C, located six residues upstream of E634C, introduced strain in the double-disulfide variants. Inspired by prior work with the respiratory syncytial virus fusion protein^40^, we introduced glycine residues flanking L628C [L628(GCG)] to increase flexibility and facilitate proper disulfide formation between I89C and L628C. This modified series was designated “G,” for glycine, followed by a construct number (Table 2, Fig. 2). Non-reducing SDS-PAGE analysis revealed that the incorporation of L628(GCG) in our double-disulfide variants modestly improved the yield and trimer-to-monomer ratio for constructs containing the A175C/E634C substitutions (Fig. 2A). Analysis by DSF also revealed that this modification eliminated a third thermal transition (T_m3_) (Fig. 2B).

**Figure 2.**
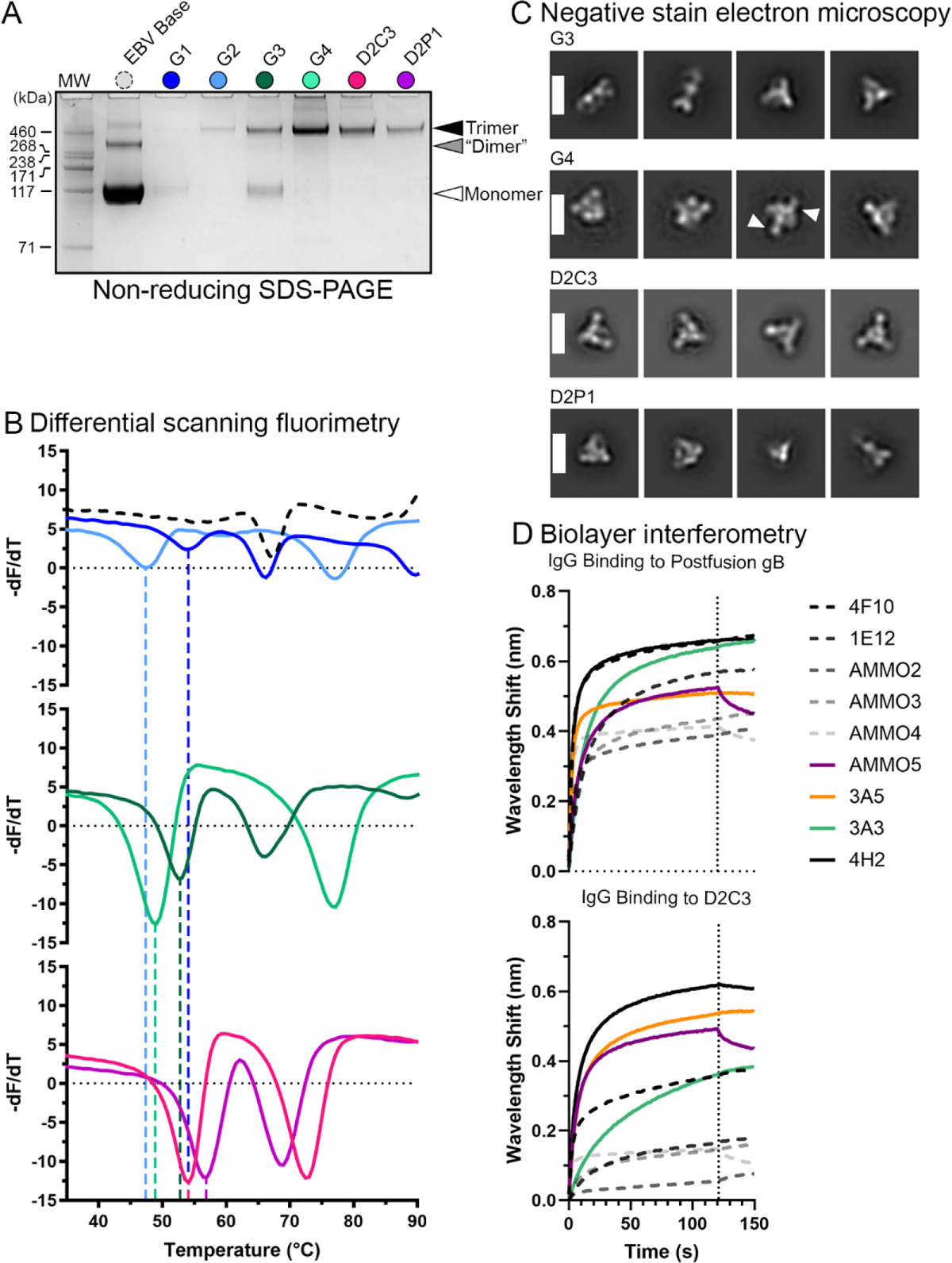
Characterization of final EBV gB combination variants. (A) Non-reducing SDS-PAGE analysis of gB variants. (B) DSF analysis of gB variant thermostability, colored as in (A). (C) nsEM 2D averages of top gB variants. White arrows highlight a splayed-out DI. Scale bars: 130 Å. (D) BLI binding of anti-gB antibodies to postfusion gB (top panel) and D2C3 (bottom panel). Neutralizing antibodies are shown as solid lines colored by target domain or colored black if the epitope is unknown. Non-neutralizing antibodies are shown as dashed black or gray lines. Vertical dotted lines represent the transition from association of IgG to dissociation. BLI data represents the average of two technical replicates.

**Table 2.** Characteristics of final EBV gB combination variants. All values were calculated from the non-reducing gel or the DSF profile in Fig. 2.

| Variant | Subs. 1 | Subs. 2 | Subs. 3 | Yield<br>Relative to<br>EBV Base | SDS-PAGE: Fraction<br>Monomer \| Dimer \| Trimer | DSF (°C)<br>T<sub>m1</sub> \| T<sub>m2</sub> \| T<sub>m3</sub> |
| --- | --- | --- | --- | --- | --- | --- |
| **G1** | N88C/<br>L628(GCG) | A175C/<br>E634C | NA | 0.05 | 0.95 \| 0.00 \| 0.05 | 54 \| 66 \| 90 |
| **G2** | N88C/<br>L628(GCG) | Q527C/<br>E634C | NA | 0.02 | 0.00 \| 0.00 \| 1.00 | 47 \| 77 \| NA |
| **G3** | I89C/<br>L628(GCG) | A175C/<br>E634C | NA | 0.28 | 0.44 \| 0.03 \| 0.54 | 53 \| 66 \| NA |
| **G4** | I89C/<br>L628(GCG) | Q527C/<br>E634C | NA | 0.35 | 0.00 \| 0.00 \| 1.00 | 49 \| 77 \| NA |
| **D2C3** | I89C/<br>L628(GCG) | A175C/<br>E634C | H316I/D320Q/<br>S325L | 0.25 | 0.00 \| 0.00 \| 1.00 | 54 \| 72 \| NA |
| **D2P1** | I89C/<br>L628(GCG) | A175C/<br>E634C | T292P | 0.13 | 0.00 \| 0.00 \| 1.00 | 57 \| 69 \| NA |

However, nsEM analysis of this construct (G3) revealed a subpopulation of postfusion gB (33%), coinciding with the presence of a monomer molecular weight band in non-reducing SDS-PAGE (Fig. 2C). Evaluation of G4 (I89C/L628(GCG), Q527C/E634C) revealed similar protrusions as seen for C4 (Fig. 2C, white arrows).

To further enhance stability, we applied ThermoMPNN and ProScan^33,41^, two tools developed to predict the effects of amino acid substitutions. Rational design informed by these resources identified a set of three substitutions (H316I/D320Q/S325L) and a proline substitution (T292P) that we applied to improve upon the G3 design. We termed these variants D2C3 (for 2 disulfide bonds, 3 computational substitutions) and D2P1 (for 2 disulfide bonds, 1 proline substitution). Both D2C3 and D2P1 exhibited 100% trimer formation by non-reducing SDS-PAGE (Fig. 2A, Table 2) and maintained similar thermostability profiles to G3 (Fig. 2B). Our nsEM analysis further confirmed that D2C3 exclusively adopted the prefusion conformation, whereas D2P1 contained particles consistent with a top-down view of postfusion gB (Fig. 2C). Additional thermostability assessments demonstrated that D2C3 retained its structural integrity after multiple freeze-thaw cycles and prolonged storage at both 4 °C and room temperature (Supplementary Fig. S6).

To further investigate antigen recognition of our gB constructs, we used biolayer interferometry (BLI) to assess antibody binding to EBV gB (Fig. 2D). Neutralizing rabbit antibodies 3A3 and 3A5, which target domains II and IV, respectively^17^, bound EBV Base and D2C3 similarly. Likewise, AMMO5^16^, a neutralizing human antibody predicted to target the disordered linker region between domains II and III^42^, bound both forms of gB. In contrast, non-neutralizing human antibodies AMMO2, AMMO3, AMMO4^16^ and 4F10^43^ exhibited notably reduced binding to D2C3, suggesting that they recognize postfusion-specific epitopes that become partially occluded in the prefusion conformation. While the epitope of 4H2 has not been reported, it is described as neutralizing^44^ and, like other neutralizing antibodies in our panel, did not exhibit postfusion specificity. These results further suggest that D2C3 adopts a prefusion conformation with distinct antigenicity compared to EBV Base.

### Cryo-EM Reveals the EBV gB Variant D2C3 Maintains a Prefusion Conformation

To analyze D2C3 in greater detail and provide a high-resolution structure of prefusion EBV gB, we froze grids of D2C3 and collected a cryo-EM dataset containing 3,895 micrographs. We were unable to identify 2D classes consistent with postfusion gB via initial unbiased blob-based picking, confirming the structural homogeneity of the construct. Subsequent template-based particle picking in CryoSPARC identified ∼2.2 million initial particles (Supplementary Fig. S7)^45^. After iterative refinements, a final 3.1 Å resolution 3D reconstruction with C3 symmetry was generated from ∼160,000 particles (Fig. 3, Supplementary Table S3). We built a model spanning EBV gB residues 44–684, although regions corresponding to residues 105–121, 184–202, 147–154, and 395–453 displayed poor map quality, suggesting intrinsic flexibility that precluded reliable modeling. To preserve consistency with previous numbering, the inserted GCG residues [L628(GCG)] were labeled 628_A_, 628_B_, and 628_C_, respectively.

**Figure 3.**
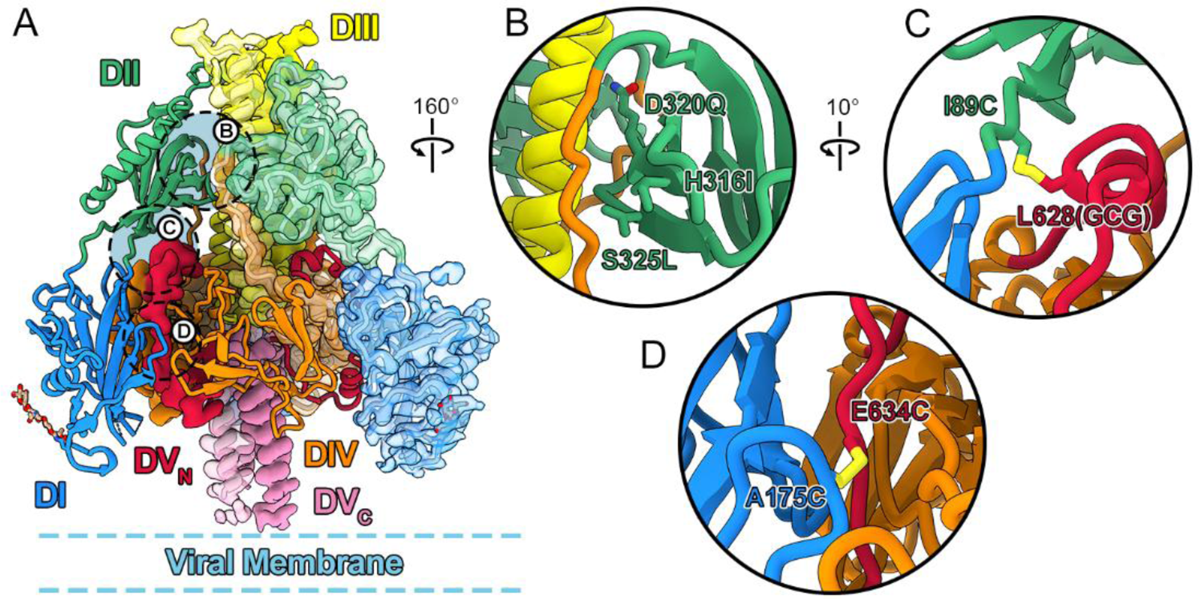
A 3.1 Å cryo-EM structure of D2C3. (A) Side view of the D2C3 model. One protomer is shown as a ribbon diagram, one protomer is shown as the opaque cryo-EM map, and one protomer is shown as the transparent cryo-EM map with the ribbon diagram of the model shown underneath. One N-linked glycan chain per protomer was built and is shown as tan sticks. The linked asparagine residue is shown as sticks colored the same as its respective ribbon diagram. (B–D) Zoomed views of substitutions that comprise D2C3. Key residues are shown as sticks. Nitrogen atoms are shown in blue, oxygen atoms are shown in red, and sulfur atoms are shown in yellow.

Comparison with the full-length prefusion HCMV gB structure^11^ revealed that D2C3 maintained a similar, compact prefusion conformation. A portion of domain I located near the membrane, containing the fusion loops (residues 112, 113, and 193–196), was disordered in the D2C3 map, likely due to the absence of the membrane-proximal region in the construct, into which the fusion loops embed. In comparison to the prefusion-like, soluble HCMV gB structure^29^, the most notable difference was in the relative location of DI, which packed more tightly into the globular core of D2C3 (Supplementary Fig. S8, black arrow), indicating better stabilization of D2C3 compared to prefusion-like, soluble HCMV gB. The D2C3 structure aligned well with the AlphaFold2 model of prefusion EBV gB, except for a one-residue register shift caused by the model’s failure to predict the α-helix preceding the L628(GCG) substitution in domain V_N_ (Supplementary Fig. S9).

To further assess the structural basis of D2C3 stabilization, we examined the cryo-EM map around the engineered disulfides and stabilizing substitutions. The I89C/L628(GCG) disulfide bond was only discernible at low-stringency contour thresholds, suggesting partial occupancy, structural heterogeneity, or radiation damage from the electron beam (Supplementary Fig. S10A-C). However, biochemical validation by non-reducing SDS-PAGE and DSF confirmed that I89C/L628(GCG) is essential for trimer formation and the thermostability of prefusion gB, even in the presence of the ThermoMPNN-derived substitutions (Supplementary Fig. S10D-E). In contrast, the A175C/E634C disulfide exhibited well-defined map features, consistent with disulfide bond formation (Supplementary Fig. S10F).

We also evaluated the map-to-model fit for the three ThermoMPNN-informed substitutions and found that each substitution was well supported by the map (Supplementary Fig. S10G-H). As expected, the H316I and S325L mutations in domain II increase hydrophobic packing interactions at the interface between domains II, III and IV. The D320Q mutation in domain II enhances intra- and interprotomer polar interactions with domain III, improving charge balance at interface between domains II and III. Taken together, these findings underscore the role of engineered disulfide bonds between domains I and II to domain V, as well as computationally predicted substitutions at the interface of domains II, III, and IV, in stabilizing prefusion EBV gB.

### Murine Immunization with Prefusion gB Modestly Enhances Neutralization Titers

To assess the immunogenicity of prefusion-stabilized EBV gB variants, three groups of five naïve BALB/c mice were immunized at days 0 and 21 with 2 µg of either postfusion gB, G3 (mixed pre- and postfusion), or D2C3 (prefusion), formulated with AddaS03 adjuvant (Fig. 4A). Sera were collected on days 21 (post-dose 1) and 35 (post-dose 2), and antibody responses were evaluated by Luminex assays. Luminex assay results suggest that immunization with constructs containing a higher proportion of prefusion gB (D2C3 > G3 > postfusion gB) was associated with an improved ratio of prefusion gB to postfusion gB binding (Fig. 4B). This trend was evident at day 21 and became more pronounced by day 35.

**Figure 4.**
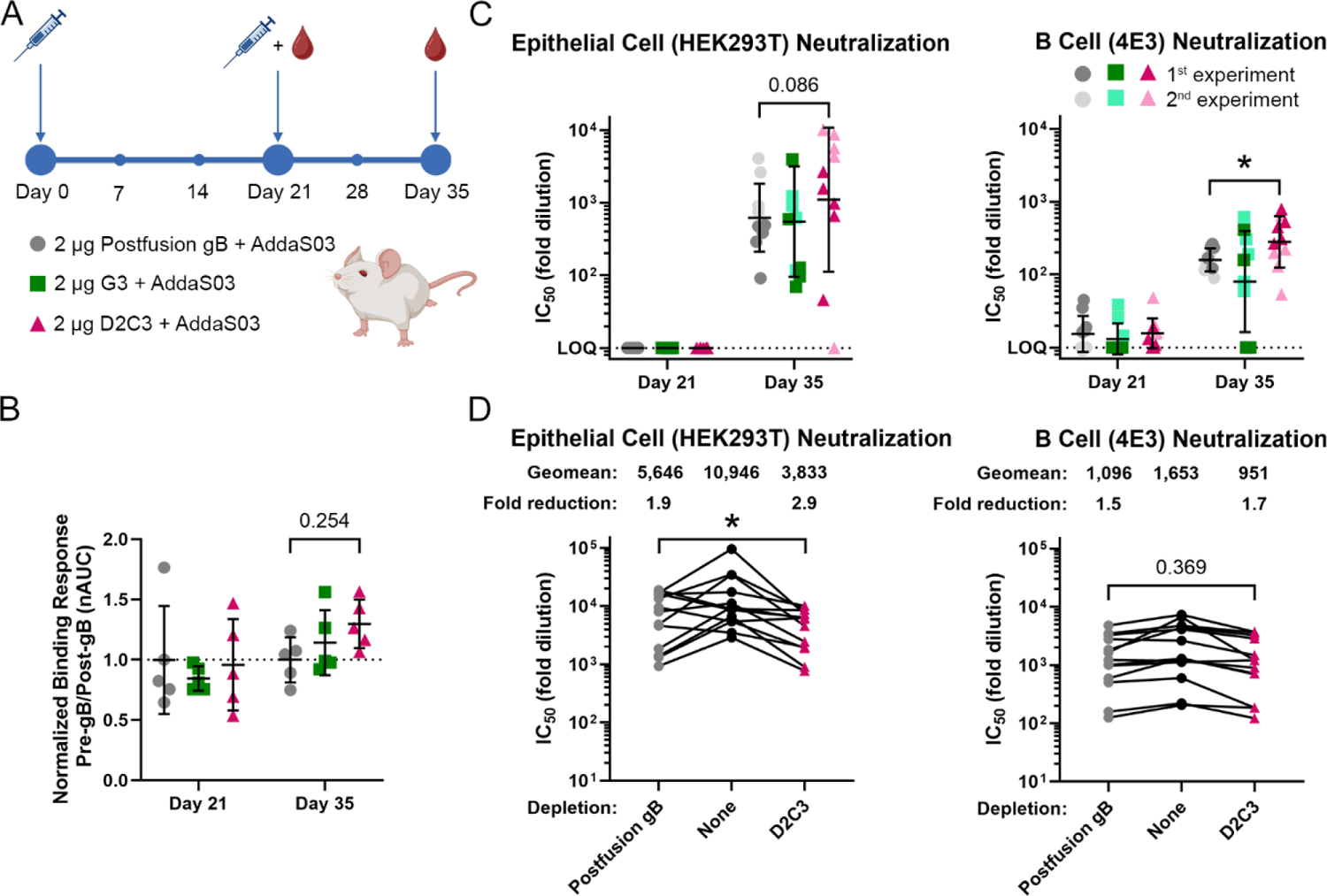
Immunogenicity of D2C3. (A) Schematic of mouse immunization study and serum collection timeline. Female BALB/c mice (7–8 weeks old, n= 5/group/experiment, two independent experiments) were immunized with one of the antigens indicated in the diagram. (B) Binding response presented as the ratio of binding to prefusion gB (Pre-gB; D2C3) divided by the binding to postfusion gB (Post-gB), normalized to the average ratio of binding observed in sera from mice vaccinated with postfusion gB. Binding was measured by a Luminex assay. Data are plotted as the average of two technical replicates. The mean is shown as a horizontal black line and error bars represent the standard deviation. (C) Neutralization by sera from immunized mice for EBV infection of the indicated cells. Data are plotted for individual mice. The geometric mean is shown as a horizontal black line and the error bars represent the geometric standard deviation. (E) Neutralization by sera from human volunteers. Sera were depleted as indicated on the x-axis. Statistical significance was determined by two-way ANOVA followed by Tukey’s HSD test (B, C) or by paired t-tests (D) performed in GraphPad Prism v10.4.2: *P < 0.05.

To assess whether prefusion gB immunization enhances neutralizing titers relative to immunization with postfusion gB, we performed live-virus neutralization assays in HEK293T epithelial cells and Akata-4E3 Burkitt lymphoma cells, an EBV-negative immortalized B cell line susceptible to EBV infection (Fig. 4C)^46,47^. After the first dose, neutralizing activity against EBV infection of epithelial cells was below the limit of quantification in all immunization groups. Following the second dose, sera from mice immunized with D2C3 exhibited a 2.0- and 1.8-fold higher geometric mean IC₅₀ than sera from mice immunized with G3 or postfusion gB, respectively. For B cells, neutralization was low after the first dose across all gB groups. Following the second dose, neutralizing titers increased in all groups, with sera from mice immunized with D2C3 exhibiting a 1.8-fold higher geometric mean IC₅₀ than sera from postfusion gB-immunized mice. Neutralization potency of sera from mice immunized with G3 remained comparable to that of mice immunized with postfusion gB. While the observed differences in neutralization of B cell infection reached statistical significance, the study was not prospectively powered, and the limited sample size and diversity may constrain the generalizability of this finding. Nonetheless, the data suggest a general trend toward moderately improved neutralizing activity following prefusion gB immunization.

Finally, we performed depletion studies using human sera from healthy, EBV-seropositive male donors 32–65 years of age to characterize the antibody response resulting from natural EBV infection (Fig. 4D; Supplementary Table S4). Sera were incubated with D2C3 or postfusion gB prior to neutralization assays. Depletion with D2C3 resulted in a 1.5- and 1.1-fold greater reduction in geometric mean IC₅₀ for epithelial and B cell neutralization, respectively, compared to depletion with postfusion gB. Although differences in epithelial cell neutralization post-depletion reached statistical significance, the modest magnitude of the effect limits its biological interpretation. Regardless, both depletion assays support a consistent trend toward improved antigenic properties for prefusion gB. This result reinforces the importance of gB as a vaccine antigen for protection against EBV infection and suggests that the human immune response to EBV can elicit prefusion gB-specific antibodies.

## Discussion

EBV infection is associated with increased risks for various cancers and multiple sclerosis, yet no licensed vaccines or therapeutics are currently available. To address this need, we leveraged rational and computational design to develop a thermostable prefusion EBV gB antigen, as confirmed by cryo-EM analysis (Fig. 3). Sera from mice immunized with prefusion EBV gB exhibited a trend toward higher neutralizing activity against EBV infection in both epithelial and B cells. Additionally, antibody depletion by prefusion gB reduced the ability of human sera from EBV-seropositive individuals to neutralize infection compared to depletion with postfusion gB (Fig. 4).

Building on our prior work demonstrating that interprotomer disulfide bonds stabilize a prefusion-like HCMV gB antigen^29^, we show here that a similar approach effectively stabilizes prefusion EBV gB by covalently linking domains I and II to domain V, though notable adjustments and additional substitutions were necessary (Fig. 2, 3). Concurrently, another group stabilized prefusion EBV gB by inserting a modified I53-50A trimerization domain into the fusion loops of domain I, along with deletions and computationally predicted substitutions to prevent refolding^48^. Together, these studies suggest three key requirements for prefusion gB stabilization. First, domain I must be locked near the globular core of gB.

This is achieved through the H222C/E657C substitutions in HCMV gB and the similar A175C/E634C substitutions in EBV gB. In the concurrent study, the engineered I53-50A domain similarly constrained domain I, enforcing comparable conformational restraints as our disulfide bond. Second, domain V must be prevented from transitioning to its extended postfusion conformation. This was accomplished through the V134C/I653C and H222C/E657C substitutions in HCMV gB and the related I89C/L628(GCG) and A175C/E634C substitutions in EBV gB. In the concurrent study, domain V stabilization was instead achieved using a Q527C/E634C substitution––an approach we also identified in our earlier designs (Fig. 1, 2). Finally, our data suggest that maintaining the proximity between domains II and III is necessary, as achieved by the H316I/D320Q/S325L substitutions identified by ThermoMPNN^33^. Although the concurrent study did not directly stabilize this interaction, deletion of the flexible linker between domains II and III likely reduced dissociation of these domains. Alternatively, their A293P substitution may rigidify the ‘elbow’ between domains I and II, thereby indirectly stabilizing the domain II/III interaction. Notably, HCMV gB has a naturally shorter domain II/III linker (Supplementary Fig. S2), potentially explaining why additional stabilization was unnecessary for that antigen. However, the splayed conformation of domain I in the prefusion-like HCMV gB antigen suggests that further stabilization of domain II/III could improve overall rigidity. While these three studies demonstrate convergent trends for gB stabilization, additional work is needed to confirm the universality of these methods in other class III fusion proteins.

While these studies outline key principles for stabilizing prefusion gB, whether this translates into improved immunogenicity across *Herpesviridae* remains uncertain. In our neutralization assays, sera from prefusion gB-immunized mice showed a consistent trend toward higher neutralizing activity (Fig. 4). However, the moderate effect size of these findings underscores the need for further studies optimizing dosing, adjuvant selection, and delivery modality to determine whether immunization with prefusion gB offers advantages over postfusion or unstabilized gB. Moreover, the reduction in binding of non-neutralizing antibodies to prefusion gB (Fig. 2D, E), coupled with the trend toward enhanced neutralization, supports further evaluation of prefusion gB as an EBV vaccine antigen. In contrast, prior studies with HCMV gB found that postfusion gB elicited equivalent neutralizing antibody titers compared to prefusion-like gB^29^. This discrepancy may be due to the increased oligomerization state (rosette formation) of postfusion HCMV gB relative to prefusion HCMV gB, as oligomerization is known to enhance immunogenicity and often increases neutralization titers^49–52^.

A recent study leveraged oligomerization to enhance EBV gB immunogenicity, showing that a multimeric postfusion gB nanoparticle elicited higher anti-gB titers as well as proportionally higher neutralization titers compared to trimeric postfusion gB^53^. However, in our EBV gB construct, we intentionally reduced oligomerization to boost expression by substituting the hydrophobic fusion loops with the less hydrophobic HSV-2 fusion loops, thereby limiting postfusion gB rosette formation^35^.

Additionally, the splayed-out conformation of domain I in prefusion-like HCMV gB may obscure or alter key neutralizing epitopes, thereby reducing the elicitation of neutralizing antibodies. These epitopes may be more accessible in the prefusion state observed for membrane-anchored HCMV gB, which more closely resembles our prefusion EBV gB antigen^11^. Consistent with this, our study found that depletion with prefusion EBV gB reduced the ability of human sera to neutralize infection compared to postfusion gB (Fig. 4D). These findings further suggest that EBV gB may be more effectively stabilized in its prefusion conformation than HCMV gB, potentially explaining its ability to elicit an improved functional antibody response. However, there may be other explanations, including intrinsic differences in the immunogenicity of the two viral proteins.

Taken together, our results support prefusion stabilization of gB as a promising vaccine development strategy for EBV. Our prefusion antigen, D2C3, exhibits improved structural stability, preferentially depletes neutralizing antibodies from human sera, and modestly enhances neutralizing antibody titers in mice compared to postfusion gB. However, given the complexity of EBV infection, an effective vaccine will likely require multiple antigens. Prior studies suggest that incorporating additional glycoproteins alongside gB, such as gH/gL, gp42, and gp350, could provide broader protection against both B cell and epithelial infection^54–59^. Furthermore, our prefusion gB construct should be a valuable tool for dissecting the antibody response to gB elicited by natural infection and vaccination. Future studies optimizing adjuvant selection, immunization regimens, and antigen combinations will be critical to fully realize the potential of prefusion gB-based vaccines in mitigating EBV-associated cancers, multiple sclerosis, and other EBV-driven diseases.

## Acknowledgements

We thank members of the McLellan Laboratory for providing helpful comments on the manuscript. We thank Dr. Axel Brilot and Dr. Evan Schwartz at the Sauer Structural Biology Laboratory at UT Austin for assistance with cryo-EM data collection and we thank Dr. Xun Zhan at the Texas Materials Institute Electron Microscopy Facility for assistance with nsEM data collection. We acknowledge the University of Texas College of Natural Sciences and award RR160023 of the Cancer Prevention and Research Institute of Texas for support of the EM facility at the University of Texas at Austin. The authors thank James Guerra, Dr. Kaci Erwin, and Dr. Mahtab Beikzadeh for technical assistance with mammalian cell culture and transfection and we thank Justine Meccio for administrative support. Funding for these studies was provided by Vaccine Company Inc. Figure 4A was created in part with BioRender.com.

## Methods

### Alignment of EBV gB, HCMV gB, and HSV-2 gB

Multiple sequence alignment was performed using Clustal Omega v1.2.4^36^ and was formatted using ESPript 3.0^60^. Amino acid sequences for EBV gB (M81 strain, GenBank: AWG92945.1), HCMV gB (Towne strain, GenBank: P13201.1), and HSV-2 gB (HG52 strain, GenBank: YP_009137179.1) were retrieved from the NCBI GenBank database. Secondary structure annotations were derived from the postfusion structure of EBV gB (PDB: 3FVC)^6^.

### Template-Based AlphaFold2 Model of Prefusion EBV gB

We generated a structural model of prefusion EBV gB using AlphaFold2 as implemented in ColabFold^32^. The full-length, prefusion HCMV gB structure (PDB: 7KDP)^11^ was used as a template.

### Design Scheme for Prefusion-Stabilized EBV gB Variants

The EBV base construct consisted of ectodomain residues 1–688 from the M81 strain (GenBank: AWG92945.1). HSV-2 fusion loop residues HR^177–178^ and RVEA^258–261^ were substituted for the EBV fusion loop residues WY^112–113^ and WLIW^193–196^, as previously described^6^. The furin cleavage site (RRRRR^428–432^) was replaced with GGSGG to prevent proteolysis. This construct was cloned upstream of a C-terminal T4 fibritin (foldon) trimerization motif, a human rhinovirus 3C (HRV3C) protease cleavage site, an octa-histidine (8xHis) tag, and a Twin-Strep tag II. Plasmids were synthesized by GenScript and cloned into the pcDNA3.1(+) expression vector with ampicillin resistance.

### Production of EBV gB Proteins for Expression Tests, SDS-PAGE, Differential Scanning Fluorimetry, and Structural Studies

Plasmids encoding EBV gB variants were transiently transfected into 80 mL FreeStyle 293-F cells (Thermo Fisher Scientific) using polyethyleneimine (PEI) and 50 µg of plasmid. Furin cleavage was evaluated for a subset of constructs by co-transfecting with 12.5 µg of plasmid encoding for the furin protease. Cultures were incubated at 37 °C, 8% CO₂, and medium was harvested after 3–6 days by centrifugation at 6,000 × g. Supernatants were 0.22 µm filtered and purified using Strep-Tactin XT 4Flow resin (IBA Lifesciences) by gravity flow. Columns (∼1 mL bed volume) were washed with 1× phosphate-buffered saline (PBS) pH 7.4, and bound protein was eluted with BXT elution buffer (IBA Lifesciences).

Eluates were analyzed by non-reducing SDS-PAGE using LDS Dye (Invitrogen Novex) and NuPAGE™ 4-12% Bis-Tris Gels (Invitrogen) run at 160 V for 40-60 minutes in 1× MES SDS Running Buffer Invitrogen). The HiMark™ Pre-Stained Protein Standard (Invitrogen) was loaded as a standard. Gels were stained with InstantBlue Coomassie Protein Stain (Abcam). Gel images were captured using a GelDoc Go imaging system (Bio-Rad).

Relative protein yield was quantified using FIJI^61^. For quantification, an identical rectangular region of interest (ROI) was applied across all lanes, encompassing the monomer, dimer, and trimer bands. The integrated density of each lane was plotted against distance, and the area under the curve (AUC) was calculated for each band using the “magic wand” tool. Relative yield was normalized to EBV Base (set to 1.00). Band ratios were determined by dividing individual band AUCs by the total AUC for all three bands within a lane.

Samples were concentrated using Amicon Ultra centrifugal filters (Millipore Sigma, 10–100 kDa cutoff) before differential scanning fluorimetry (DSF) analysis. Following DSF, samples were flash-frozen in liquid nitrogen and stored at −80 ° C.

### Differential Scanning Fluorimetry

For DSF experiments, EBV gB variants were diluted to 0.684 mg/mL in 20 mM Tris (pH 8.0), 200 mM NaCl, 2 mM CaCl₂, and 0.02% (w/v) sodium azide (SEC Buffer). Samples were prepared in triplicate, with 15 µL per well in a white-walled 96-well PCR plate (VWR).

SYPRO Orange dye was prepared by diluting 5,000× SYPRO Orange Protein Gel Stain in DMSO (Thermo Fisher Scientific) to a 66× stock solution and 5 µL of dye solution was added to each well, yielding final concentrations of 0.513 mg/mL protein and 16.5× dye. Plates were sealed, centrifuged at 1,000 × rcf for 2 min, and loaded into a LightCycler 480 differential scanning fluorimeter (Roche) equipped with a 100 W xenon excitation lamp, as previously described^29^. DSF was performed using a 465 nm excitation filter (25 nm half bandwidth) and a 580 nm emission filter (20 nm half bandwidth) while gradually increasing the temperature from 25 °C to 90 °C.

Fluorescence was monitored in the LightCycler 480 software, and thermal denaturation curves were plotted as the change in fluorescence with respect to temperature (-dF/dT) before reformatting in GraphPad Prism v10 (Dotmatics).

### Negative-Stain Electron Microscopy

Purified EBV gB constructs were centrifuged at >20,000 × g for 7–10 min and diluted to 0.05 mg/mL in 20 mM Tris (pH 8.0), 200 mM NaCl, and 0.02% (w/v) sodium azide. Copper-supported carbon grids (400-mesh Formvar) were glow discharged by an EM ACE600 high vacuum sputter coater (Leica Microsystems) for 30 s at 10 mAmps. Diluted samples were immediately applied to glow-discharged grids and stained with 0.22 µm-filtered 2% uranyl acetate (w/v). Once air-dried, grids were loaded onto a Japan Electron Optics Laboratory (JEOL) NEOARM electron microscope equipped with a OneView camera (Gatan) and operated at 200 kV with a nominal magnification of 50,000×. The C1 aperture was inserted, and micrographs were acquired in 2k × 2k mode (pixel size: 4.32 Å). Micrographs were saved as .dm3 files using Digital Micrograph (Gatan) before conversion to .mrc format using e2proc2d.py (EMAN2)^62^. Converted files were imported into CryoSPARC (Structura Biotechnology) for contrast transfer function (CTF) correction, particle picking, and 2D classification^45^.

### Cryo-EM Sample Preparation and Data Collection

Flash-frozen EBV gB variants were thawed at 37 °C, centrifuged at >20,000 × g for 7–10 min, and immediately injected onto a Superose 6 Increase 10/300 GL column (Cytiva). Size-exclusion chromatography (SEC) was performed in SEC buffer. Desired fractions were pooled and concentrated.

UltrAuFoil 1.2/1.3 grids (Electron Microscopy Sciences) were glow-discharged for 30 s at 25 mAmps (PELCO easiGlow™ Glow Discharge Cleaning System). EBV gB variants were diluted to 6.2 mg/mL in SEC buffer with 2 mM CHAPS (0.25× CMC or 0.123% w/v), which was added immediately before 4 µL of sample was applied to glow-discharged grids.

Grids were plunge-frozen in liquid ethane using a Vitrobot Mark IV (Thermo Fisher Scientific) set to 100% humidity and 4 °C, with a blot time of 6 s, blot force of 8 s, and wait time of 0 s. Cryo-EM datasets were collected at 150,000× magnification, corresponding to a calibrated pixel size of 0.933 Å, using an FEI Glacios electron microscope (Thermo Fisher Scientific) operating at 200 kV and equipped with a Falcon 4 direct electron detector (Thermo Fisher Scientific). SerialEM v3.9.0 beta^63^ was used to acquire 4,541 total exposures at 0° tilt for the D2C3 dataset.

### Cryo-EM Data Processing, Model Building, and Refinement

Gain reference correction was applied before importing micrographs into CryoSPARC Live (Supplementary Fig. 7)^45^. Motion correction, patch CTF estimation, defocus estimation, micrograph curation, and initial particle picking were performed in CryoSPARC Live. Of the 4,541 collected micrographs, 3,895 were accepted for further processing.

Micrographs and selected particles were exported into CryoSPARC v4.5.3 for *ab initio* refinement, where the resulting volume was used for template-based particle picking. Particles underwent 2D classification, *ab initio* refinement, heterogeneous refinement, homogeneous refinement, and final non-uniform refinement. Automatic post-processing was performed using the high-resolution option in DeepEMhancer with half maps, but not a mask, supplied as inputs^64^.

Our AlphaFold2-predicted prefusion EBV gB model was used as the initial model for D2C3. The model was rigid-body fitted into the refined map using ChimeraX^65^, followed by iterative refinement in PHENIX^66^, COOT^67^, and ISOLDE^68^. Structural figures were generated using ChimeraX^65^.

### Antibody Binding Assay

To prepare antigen conjugated beads, MagPlex®-Avidin microspheres (Diasorin) designed for postfusion gB (postfusion trimer, region 18), variant G3 (pre- and postfusion trimer, region 37), variant D2C3 (prefusion trimer, region 39), irrelevant antigen (MERS-CoV S-protein 3P.GCN4^69^, region 13) and unconjugated blank beads (Region 12), were counted using a Countess S3 cell counter (Thermo Fisher) and collected via magnetic separation. Storage buffer was removed and beads were resuspended in 3 mL PBS pH 7.4 + 1% bovine serum albumin (BSA, w/v), and 5 µg antigen/100 kDa antigen/1 × 10^6^ beads was added. After 2 h, beads were washed and resuspended in 1 mL Luminex wash buffer [0.1% BSA (w/v), 0.02% Tween 20 (w/v) in PBS pH 7.4]. The post-conjugation antigenicity of each antigen was validated by measuring binding to monoclonal antibodies 3A3 and 3A5 for gB-conjugated beads^17^ or JC57-14 for the irrelevant control MERS antigen^70^. All monoclonal antibodies for antigenicity validation were purchased from GenScript. To assess antibody binding, murine serum samples were diluted 75-fold, followed by an 8-point 5-fold serial dilution series in Luminex assay diluent [1% non-fat milk (w/v), 5% FBS (w/v), 0.05% Tween 20 (w/v) in PBS pH 7.4]. 25 µL of diluted serum was added to 25 µL of antigen-conjugated microsphere suspension (at least 1000 microspheres/antigen). After 30 min, microspheres were pelleted with a magnetic separator and washed three times with 300 µL of Luminex wash buffer.

Microspheres were incubated with 100 µL of goat anti-mouse phycoerythrin-labelled IgG (2 µg/mL, SouthernBiotech) for 30 min, pelleted with a magnetic separator, washed three times with 300 µL of Luminex wash buffer, and resuspended in 200 µL of Luminex wash buffer for analysis. Mean fluorescence intensity (MFI) was measured using a Luminex xMAP Intelliflex. MFI values were background-corrected by subtracting signals from unconjugated blank beads. Area under the MFI curve (AUC) was calculated using GraphPad Prism 10.4.

### Viral Production

Live EBV containing a green fluorescent protein (Akata-EBV GFP) reporter was isolated from mutant Akata BX-1 cells^46,47^, which contain a recombinant EBV with GFP/Neomycin gene placed in the BXLF1 region of the viral genome. Cells were seeded at a density of 0.3–1.0 × 10^6^ cells/mL in RPMI-1640, 10% heat inactivated FBS (w/v), 1× GlutaMAX supplement (Gibco, 35050061), 100 U/mL Penicillin-Streptromycin (R10 media) + 350 µg/mL G418 (Corning, 30-234-CI) at 37 °C, 5% CO_2_ in approximately 1 L. Once expanded, cells were resuspended to 4.0 × 10^6^ cells/mL in RPMI-1640 + 1% heat inactivated FBS (w/v) + 1× GlutaMAX supplement (R1 media) and virus was induced by treatment with 100 µg/mL goat anti-human IgG F(ab’)_2_ (MP Biomedicals, 0855049). Cells were incubated for 4 h and subsequently diluted in R1 media to a final concentration of 2.0 × 10^6^ cells/mL. Cells were incubated for 5 d and then pelleted by centrifugation at 5,000 rpm for 10 min at 4 °C. Supernatant was collected, 0.8 µm cellulose nitrate filtered (Thermo Fisher), and treated with 100 µg/mL bacitracin. Virions were then isolated by centrifugation at 21,000 × *g* for 90 min at 4 °C using a fixed angle rotor. Supernatant was decanted and virus was resuspended at 100-fold concentration in RPMI-1640 + 100 µg/mL bacitracin; aliquots were stored in the vapor phase of liquid N_2_ until use.

### Viral Culture and Neutralization Assays

Virus titer and neutralization were performed based on previously described methods^46^. Briefly, Akata-EBV-GFP virus was titrated on 4E3^71^ and HEK293T (ATCC, CRL-3216) target cell lines. Virus was serially 2-fold diluted in R10 media without phenol red starting from a 1:5 starting dilution and 30 µL/well of virus (or R10 as an untreated control) was added to a 96-well black wall, transparent bottom plate (Corning, 3904). 30 µL/well of R10 media was added to each well and 4.4 × 10^4^ 4E3 cells/well or 2.0 × 10^4^ HEK-293T cells/well in 60 µL was added to each well and incubated for 3 d at 37 °C, 5% CO_2_. Cells were analyzed by high content imaging as described below, and the dilution of virus needed to target a 20% infectivity was determined for each virus lot.

For neutralization assays, all sera were heat inactivated at 56 °C for 30 min. B cell neutralization was assessed in 4E3 cells based on previously described methods^46^ and epithelial cell neutralization was assessed in HEK-293T cells. Briefly, 4E3 cells were maintained at 37 °C, 5% CO_2_ in R10 media and HEK-293T cells were maintained in R10 media without phenol red. For neutralization, 30 µL/well of sera (or controls) were prepared in a 96-well plate at a starting dilution of 1:5 in R10 and serially 5-fold diluted in R10. Then, 30 µL/well of EBV-GFP virus was added to each well to obtain a top dilution of 1:10, and sera were incubated with virus for 1h at 37 °C. Uninfected cells and virus-only treatments were prepared as positive and negative controls, respectively. 4.4 × 10^4^ 4E3 cells/well or 2.0 × 10^4^ HEK-293T cells/well in 60 µL were added to virus-containing wells and incubated for 3 d at 37 °C, 5% CO_2_.

High content image collection and analysis was conducted using an ImageXpress® Pico (Molecular Devices). Image-based neutralization assays were run in R10 media without phenol red to reduce background fluorescence. Plates were directly transferred from the 37 °C incubator 3 d after infection to the instrument and imaged using a 10× objective in a single plane. Transmitted light and GFP images were acquired at automatically determined exposure settings, with two fields of view collected per well. Data were analyzed using the Pico’s automated cell identification algorithm, which uses a transmitted light image to identify individual cells and GFP thresholds were defined based on cell-only and virus-only infection conditions. Imaging settings were maintained consistent across all assays for each cell type. Following collection of counts for GFP positive and negative cells, serum concentrations giving 50% neutralization relative to the positive and negative controls (IC_50_) were determined by 4-parameter unconstrained logistic regression using GraphPad Prism 10.4.

### Human Serum Depletion

Healthy, EBV-seropositive adult human serum samples, with no documented linkage to an EBV-mediated pathology or disease (Supplementary Table S4), were purchased from BioIVT and heat-inactivated at 56 °C for 30 minutes. Streptavidin-coated magnetic beads (Pierce, 88817) were coated with biotinylated gB antigens, or left uncoated as a mock depletion, at a density of 50 μg antigen/1 mg of beads in PBS pH 7.4 with 10% FBS (w/v) and 0.1% Tween 20 (w/v) by end-over-end mixing at 4 °C for 2 h. Coated beads were kept at 4 °C until use. To deplete antigen specific antibodies, 0.25 mg of antigen-coated beads were magnetically adhered to a 96-well plate and binding buffer removed. Beads were incubated with 50 μl of serum overnight at 4 °C with end-over-end mixing. Beads were magnetically adhered to the plate and depleted serum was removed and stored at −80 °C until use.

### Antigen Production and Purification

Postfusion gB and D2C3 (prefusion gB) containing either a Foldon-His-Strep or a GCN4-AviTag-His were produced in Expi293F cells and purified via His-tag purification as described below. Expi293F cells were cultured in Expi293 media (Gibco) at 37 °C, 120 RPM, and 5% CO_2_. For transfection, cells were grown to a density of 3 × 10^6^ cells/mL and transfected with plasmid DNA at 1 µg DNA/mL of cell culture using ExpiFectamine293 transfection reagent (Gibco) according to manufacturer recommendations. For constructs containing AviTag sequences, transfections were performed using a 1:1 ratio of gB:BirA plasmid in which BirA biotin ligase is expressed under control of a CMV promoter. Transfection in this manner facilitates in-cell biotinylation of the proteins produced. Cells were harvested 4–6 days post transfection by centrifugation at 7000 × g followed by supernatant filtration using a 0.22-µm filter.

Filtered supernatant was diluted with 10× PBS to a final concentration of 2.5× PBS (pH 7.4) and loaded onto a HisTrap excel (Cytiva) column pre-equilibrated in 2.5× PBS pH 7.4. Proteins were eluted with 250 mM imidazole in 2.5× PBS pH 7.4. Pooled HisExcel elutions were concentrated using Amicon Ultra centrifugal filters (Millipore Sigma, 100 kDa cutoff) to 0.5 mL. Concentrated proteins were filtered using a 0.22-µm spin filter before further purification on a Cytiva S6 10/300 GL (PN 29091596) or a Cytiva Superdex 200 Increase 100/300GL (PN 28990944) SEC column pre-equilibrated in 20 mM Tris pH 7.5, 150 mM NaCl. Protein-containing fractions were pooled, snap-frozen, and stored at −80 °C until use.

### Biolayer Interferometry

Biolayer interferometry analysis was performed using an Octet R8 instrument. Antibodies were procured from GenScript with human Fc domains and stored in 1× TBS pH 7.4. V_H_ and V_L_ sequences may be found in the Supporting Information files. Antibodies and purified gB proteins were diluted to 10 µg/mL in 1× Sartorius kinetics buffer [PBS pH 7.4 + 0.1% BSA (w/v), 0.02% Tween 20 (w/v), and a microbicide, Kathon] and pipetted into black-walled, black-bottom plates (200 µL/well). His1K tips were dipped into gB protein wells until an ∼0.4 nm loading threshold was reached and then subsequently dipped into antibody wells to assess binding association for 120 s. Dissociation was monitored for 30 s by moving tips into a well containing buffer alone.

### Vaccine Formulation and BALB/c Mouse Immunizations

Female BALB/c mice (age 7–8 weeks) were purchased from Charles River Laboratories and acclimated for at least one week in a contract vivarium (Pacific Immunology) prior to immunization. HisTrap-purified EBV gB antigens were diluted to 0.02 mg/mL in 5% sucrose (w/v), 20 mM Tris pH 7.5, and 150 mM NaCl and admixed 1:1 with AddaS03 (InvivoGen). Mice were injected intramuscularly with a 50 µL dose of antigen in each hind leg, to deliver 100 µL of vaccine formulation per immunization. To obtain blood via retroorbital bleeding, a capillary tube was inserted into the medial canthus. Blood was then transferred to an Eppendorf or BD Microtainer Blood Collection Tube (BD 365967), allowed to clot at room temperature for approximately 2 h, and centrifuged at 4000 rpm in a tabletop centrifuge. The serum layer was transferred to an Eppendorf tube and frozen at −80 °C. All mouse studies were conducted in accordance with IACUC-approved protocols.

### Data Processing, Artificial Intelligence and Statistical Analysis

Electron microscope processing was performed using CryoSPARC v 4.5.3 and subsequent versions^45^. Structural biology applications were compiled and configured by SBGrid^72^. ChatGPT was used to aid in manuscript editing. Data were plotted using GraphPad Prism 10.4.0 and subsequent versions.

### Data, Materials, and Software Availability

All data needed to evaluate the conclusions in the paper are present in the paper and/or the Supplementary Materials. Electron microscopy maps and structure models for prefusion gB is accessible through EMD-70288 and PDB ID: 9OAL.

## Notes

### Competing Interest Statement

The authors declare the following competing interests: A.E.P., D.J.S., J.H., H.C., S.P., J.-L.C., J.E.L., and P.A.-B.W. are employees and possible shareholders of Vaccine Company, Inc. R.S.M., C.M.A, P.O.B., M.R.S., and J.S.M. are inventors on US patent application no. 63/689,195, entitled Prefusion-stabilized EBV gB Proteins.

