## Supplementary Information for "Design, Structure, and Immunogenicity of a Soluble Prefusion-stabilized EBV gB Antigen"

**Supplemental Figures and Tables**

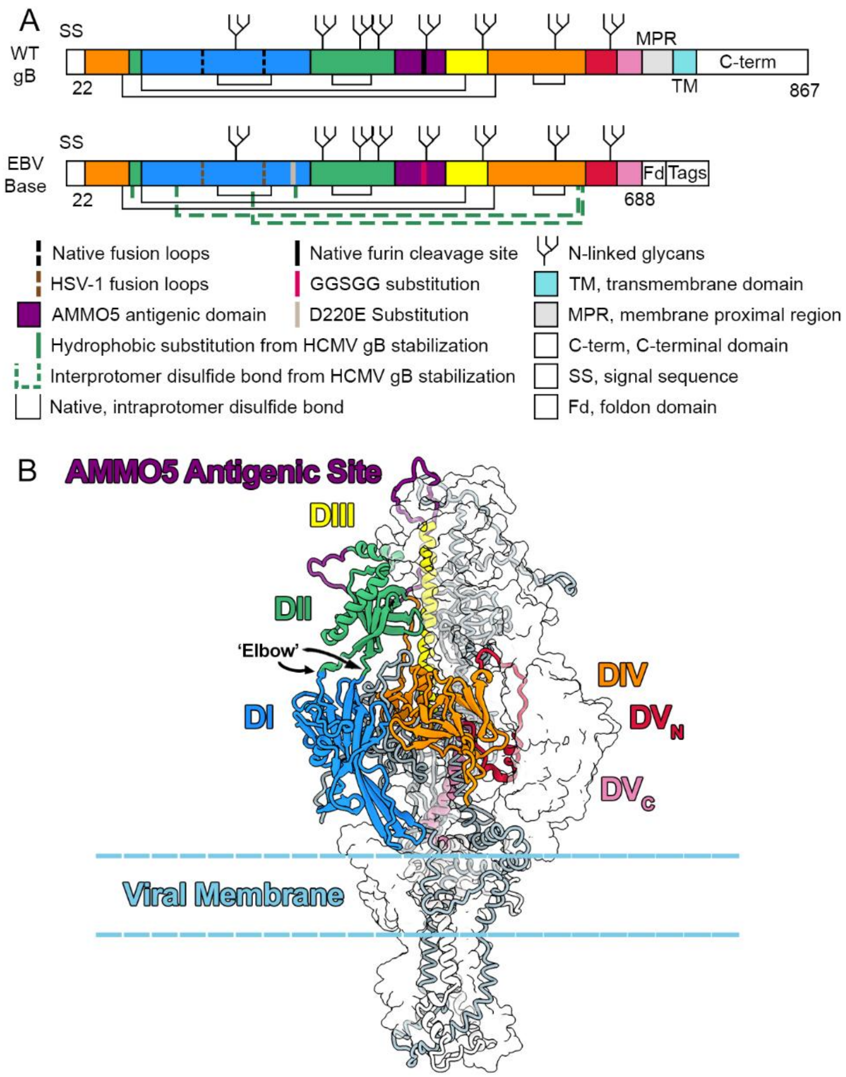


**Supplemental Figure 1 | Design theory for EBV gB Base, colored by domain.** (A) Schematic of wildtype (WT) gB and ectodomain base construct (EBV Base). EBV Base contains modifications listed in the diagram. (B) AlphaFold2 model of prefusion WT gB. Trimeric prefusion HCMV gB (PDB ID: 7KDP)[1] was used as a template. One protomer is shown as a ribbon and is colored as in (A), one is gray and shown as a trace of the α-carbon backbone, and one is shown as a transparent surface. The 'elbow' allowing DI to splay out in partially prefusion-stabilized ectodomain constructs is labeled in black and marked with arrows.

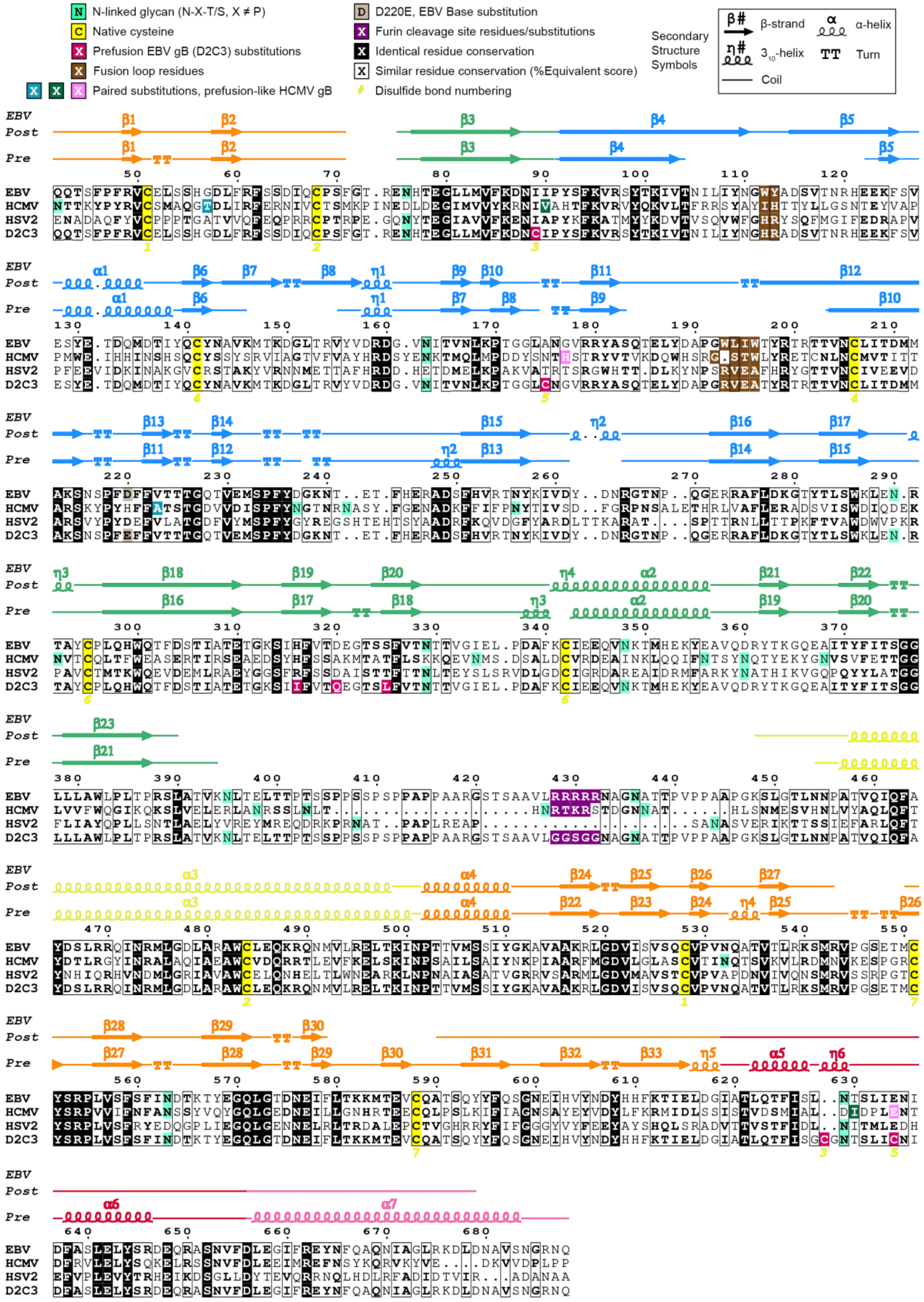
N-linked glycan (N-X-T/S, X ≠ P)
Native cysteine
Prefusion EBV gB (D2C3) substitutions
Fusion loop residues
Paired substitutions, prefusion-like HCMV gB
D220E, EBV Base substitution
Furin cleavage site residues/substitutions
Identical residue conservation
Similar residue conservation (%Equivalent score)
Disulfide bond numbering
Secondary Structure Symbols
β-strand
α-helix
3₁₀-helix
Turn
Coil
EBV Post
Pre
EBV
HCMV
HSV2
D2C3

**Supplemental Figure 2 | Sequence alignment of EBV gB, HCMV gB, HSV-2 gB, and prefusion EBV gB (D2C3).** Multiple sequence alignment of EBV gB (M81 strain, GenBank: AWG92945.1), HCMV gB (Towne strain, GenBank: P13201.1), and HSV-2 gB (HG52 strain, GenBank: YP_009137179.1) was performed using Clustal Omega v1.2.4[2] and formatted with ESPript v3.0[3]. The alignment is limited to residues 42–688, corresponding to the first and last residues resolved for the postfusion and prefusion EBV gB structures, respectively. Gaps in secondary structure correspond to unresolved residues. Select residues and features are marked as indicated in the legend above. Secondary structure elements were derived from the postfusion EBV gB crystal structure (Post-gB, PDB: 3FVC)[4] or from the prefusion EBV gB structure (Pre-gB) and are colored by domain as in Supplementary Fig. S1.

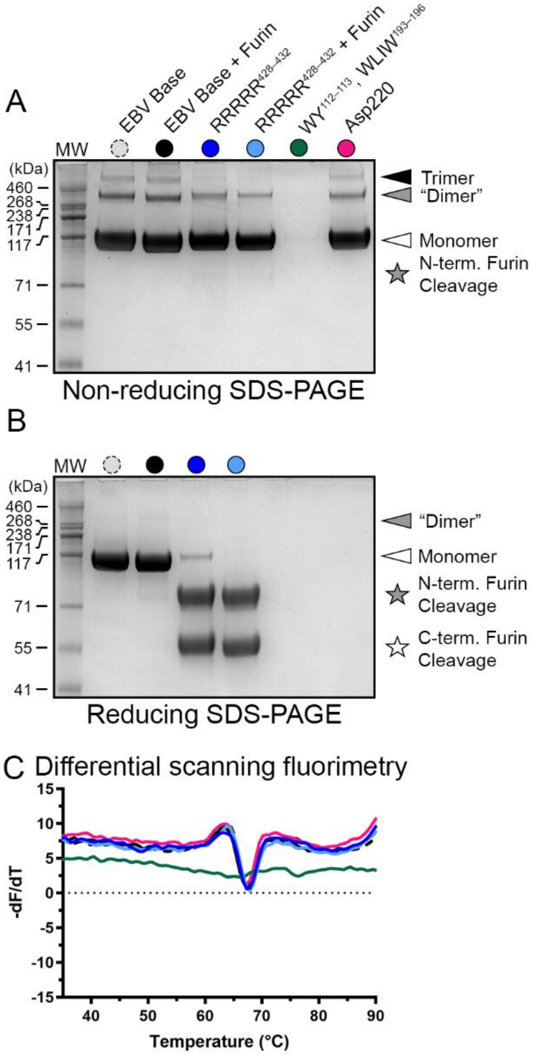


**Supplemental Figure 3 | Effects of individual modifications on EBV Base.** (A) Non-reducing SDS-PAGE analysis of gB variants. (B) Reducing SDS-PAGE analysis of gB variants. (C) DSF analysis of gB variant thermostability colored as in (A).

**Table S1 | Effects of substitutions in the EBV Base construct.** All values were calculated from the non-reducing gel or the DSF profile in Supplementary Fig. S3.

| Variant | Yield Relative to EBV Base | SDS-PAGE: Fraction Monomer \| Dimer \| Trimer | DSF (°C) $T_{m1}$ \| $T_{m2}$ |
|---|---|---|---|
| **EBV Base** | 1.00 | 0.81 \| 0.16 \| 0.03 | 68 \| NA |
| **RRRRR$^{428–432}$** | 0.99 | 0.92 \| 0.08 \| 0.00 | 67 \| NA |
| **WY$^{112–113}$, WLIW$^{193–196}$** | 0.12 | 1.00 \| 0.0 \| 0.0 | 66 \| 76 |
| **Asp220** | 1.06 | 0.88 \| 0.11 \| 0.01 | 67 \| NA |

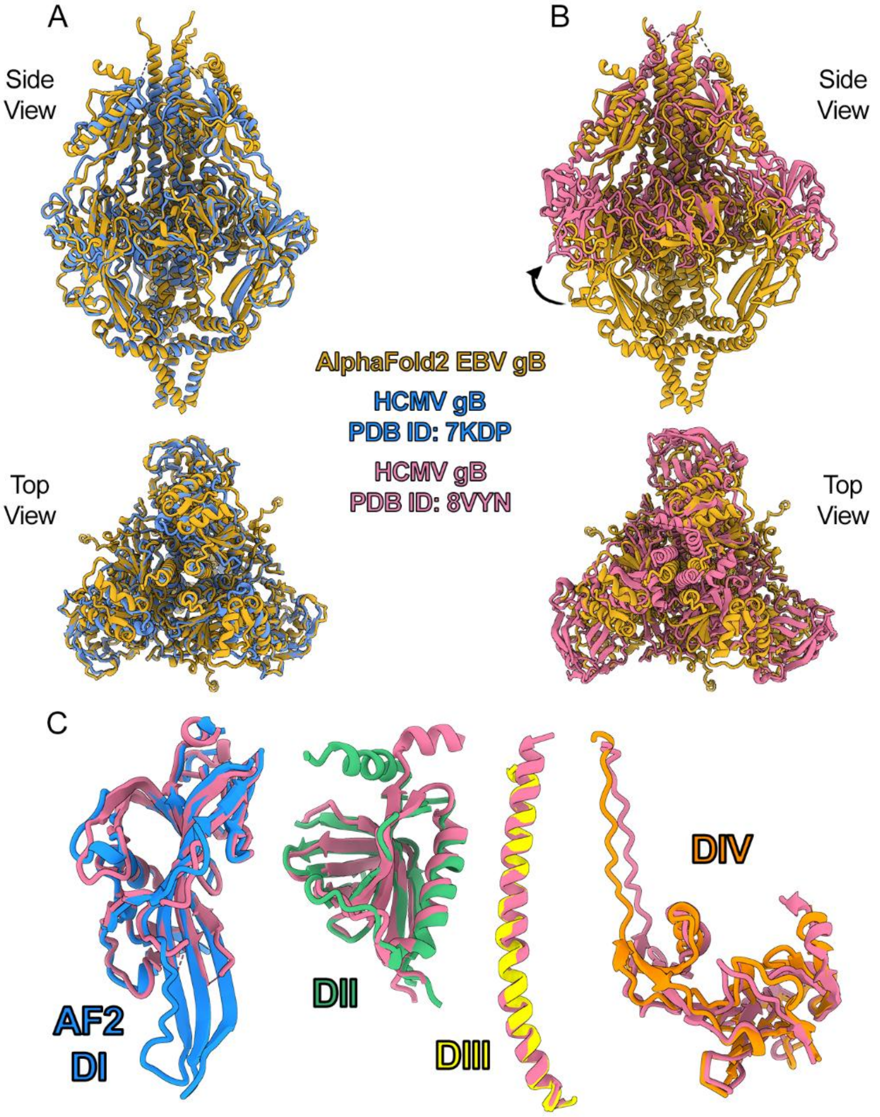


**Supplemental Figure 4 | Alignment of the AlphaFold2 model of prefusion EBV gB with two HCMV gB prefusion structures.** (A) The EBV gB prefusion model (orange, AlphaFold2)[5] aligned with membrane-anchored HCMV gB structure (blue, PDB:7KDP)[1]. (B) The EBV gB prefusion model aligned with the HCMV gB ectodomain structure (pink, PDB: 8VYM)[6]. The black arrow highlights the movement of domain I (DI) between the EBV gB model and the HCMV gB ectodomain structure. (C) Pairwise alignments of DI–DIV between the EBV gB model and the HCMV gB ectodomain structure. The EBV gB model is colored by domain as in Supplementary Fig. S1. HCMV gB is colored as in (B). All alignments were performed using the MatchMaker function in ChimeraX[7]. Individual domains were aligned based on residues within the respective domain.

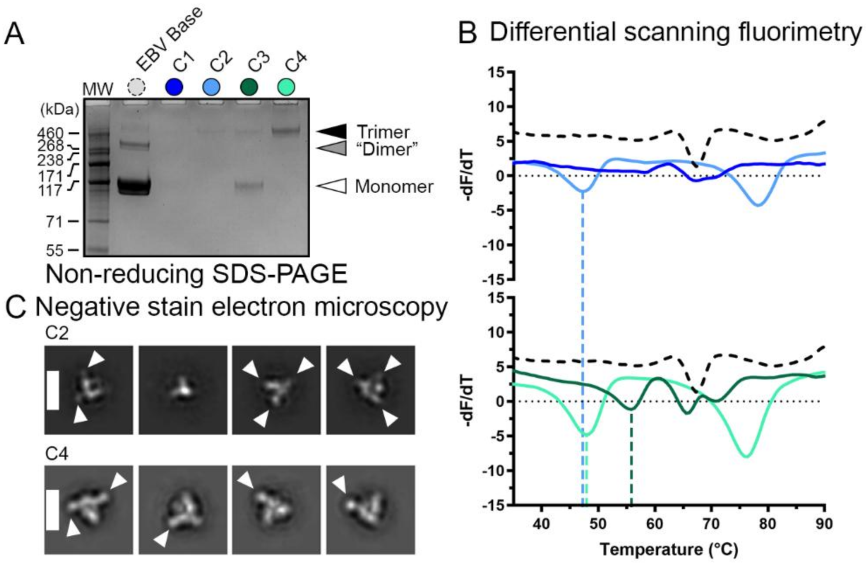


**Supplemental Figure 5 | Characterization of EBV gB combination variants.** (A) Non-reducing SDS-PAGE analysis of gB variants. (B) DSF analysis of gB variant thermostability, colored as in (A). (C) Negative-stain electron microscopy (nsEM) 2D averages of top double-disulfide variants. White arrows highlight a splayed-out DI. Scale bars: 130 Å.

**Table S2 | Characteristics of EBV gB combination variants.** All values were calculated from the non-reducing gel or the DSF profile in Supplementary Fig. S5.

| Variant | Subs. 1 | Subs. 2 | Yield Relative to EBV Base | SDS-PAGE: Fraction Monomer \| Dimer \| Trimer | DSF (°C) $T_{m1}$ \| $T_{m2}$ \| $T_{m3}$ |
|---|---|---|---|---|---|
| C1 | N88C/ L628C | A175C/ E634C | NA | NA | 67 \| NA \| NA |
| C2 | N88C/ L628C | Q527C/ E634C | 0.02 | 0.00 \| 0.00 \| 1.00 | 47 \| 78 \| NA |
| C3 | I89C/ L628C | A175C/ E634C | 0.14 | 0.73 \| 0.00 \| 0.27 | 55 \| 65 \| 71 |
| C4 | I89C/ L628C | Q527C/ E634C | 0.13 | 0.00 \| 0.00 \| 1.00 | 47 \| 76 \| NA |

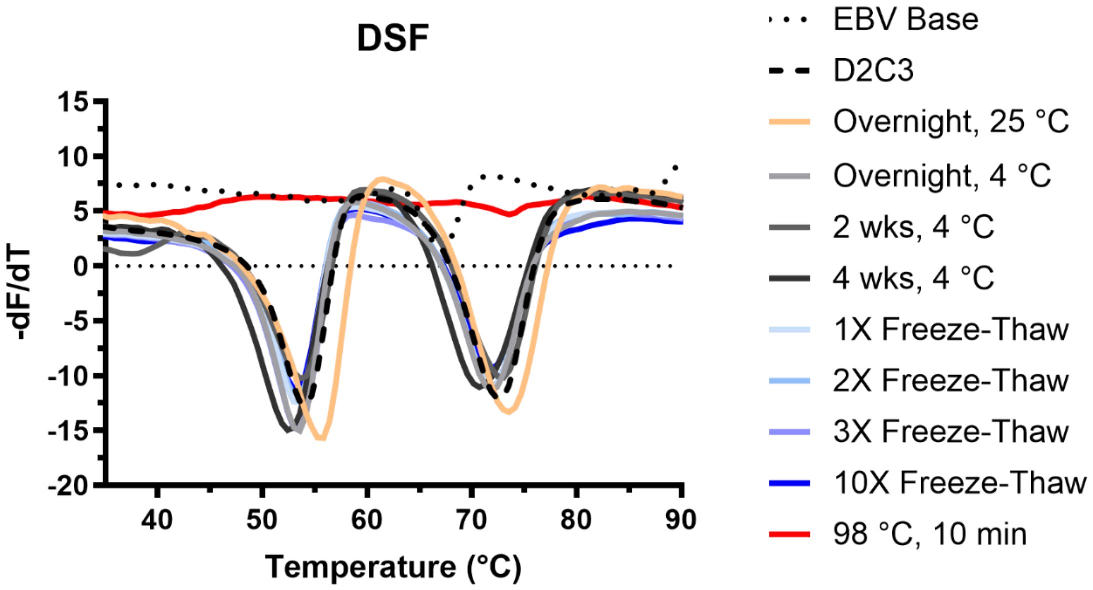


**Supplemental Figure 6 | D2C3 storage stability test.** Differential scanning fluorimetry (DSF) analysis of D2C3 after various storage conditions and lengths. For freeze-thaws, protein was flash frozen in liquid nitrogen and immediately thawed in a 37 °C water bath for the indicated number of rounds.

**Table S3 | Cryo-EM Data collection and refinement statistics.**

| **Sample** | D2C3 (prefusion EBV gB) |
|---|---|
| PDB ID | 9OAL |
| EMDB code | EMD-70288 |
| **EM data collection** | |
| Microscope | FEI Glacios |
| Voltage (kV) | 200 |
| Detector | Falcon 4 |
| Magnification (nominal) | 150,000 |
| Pixel size (Å/px) | 0.933 |
| Exposure rate (e-/pix/sec) | 9 |
| Exposure (e-/Å$^2$) | 49 |
| Decous range (μm) | 1.5–3.0 |
| Tilt angle (°) | 0 |
| Micrographs collected | 4,541 |
| Micrographs used | 3,895 |
| Patricles extracted (total) | 2,178,814 |
| Automation software | SerialEM |
| **3D reconstruction statistics** | |
| Particles | 162,928 |
| Symmetry | C3 |
| Map sharpening B-factor | -121.7 |
| Unmasked resolution at 0.5 FSC (Å) | 3.8 |
| Masked resolution at 0.5 FSC (Å) | 3.4 |
| Unmasked resolution at 0.143 FSC (Å) | 3.5 |
| Masked resolution at 0.143 FSC (Å) | 3.1 |
| **Model refinement and validation statistics** | |
| Refinement package | PHENIX |
| Composition | |
| Non-hydrogen atoms | 13,059 |
| Protein residues | 1,620 |
| Ligands | 9 (3 BMA, 6 NAG) |
| RMSD bonds (Å) | 0.005 |
| RMSD angles (°) | 0.996 |
| Average B-factors | |
| Protein resiudes | 76.8 |
| Ligands | 30.0 |
| Ramachandran | |
| Favored (%) | 96.7 |
| Allowed (%) | 3.3 |
| Outliers (%) | 0 |
| Rotamer outliers (%) | 0.21 |
| Clash score | 2.68 |
| Cβ outliers (%) | 0 |
| CaBLAM outliers (%) | 0.96 |
| MolProbity Score | 1.26 |
| EMRinger score | 3.38 |
| CC (mask) | 0.81 |

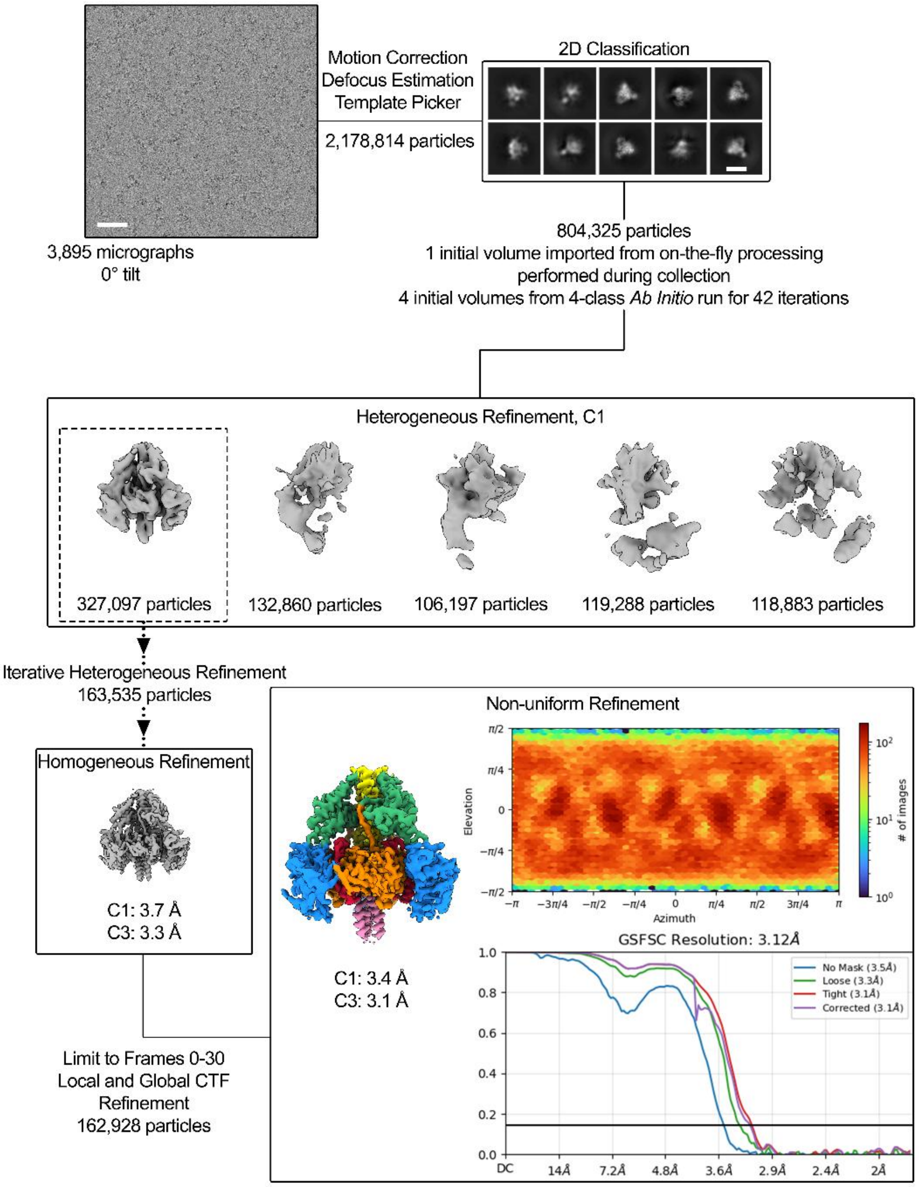


**Supplemental Figure 7 | Representative cryo-EM processing workflow for D2C3.** Scale bars are 50 nm and 10 nm for the micrograph and 2D classes, respectively. The final cryo-EM map from non-uniform refinement is colored by domain. All maps are unsharpened and the final map, sharpened by DeepEMhancer[8], may be found in Figure 3. All processing was performed in CryoSPARC v4.5.3[9] and later versions.

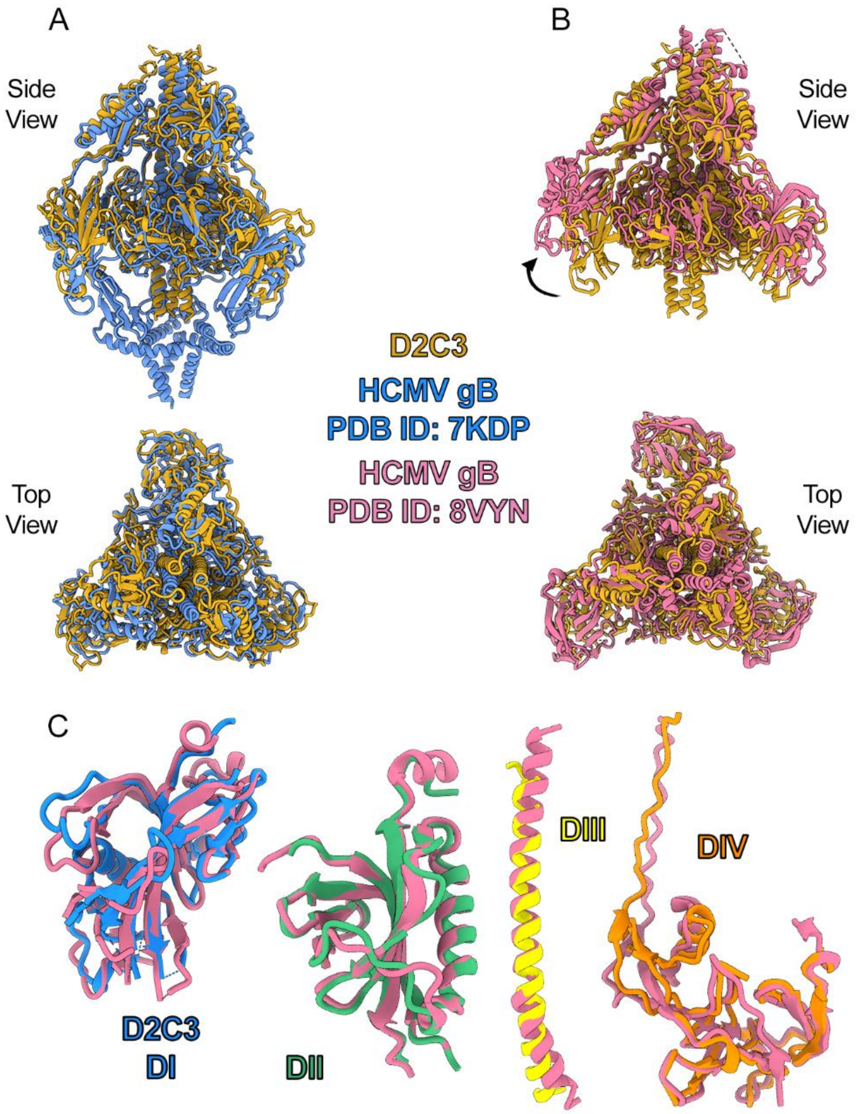


**Supplemental Figure 8 | Alignment of D2C3 structure with two HCMV gB prefusion structures**. (A) D2C3 (orange) aligned with membrane-anchored HCMV gB construct (blue, PDB:7KDP)[1]. (B) D2C3 aligned with the HCMV gB ectodomain construct (pink, PDB: 8VYM)[6]. The black arrow highlights the movement of domain I (DI) between D2C3 and the soluble HCMV gB ectodomain construct. (C) Pairwise alignments of DI–DIV between D2C3 and the HCMV gB ectodomain construct. D2C3 is colored by domain as in Supplementary Fig. S1. HCMV gB is colored as in (B). All alignments were performed using the MatchMaker function in ChimeraX[7]. Individual domains were aligned based on residues within the respective domain.

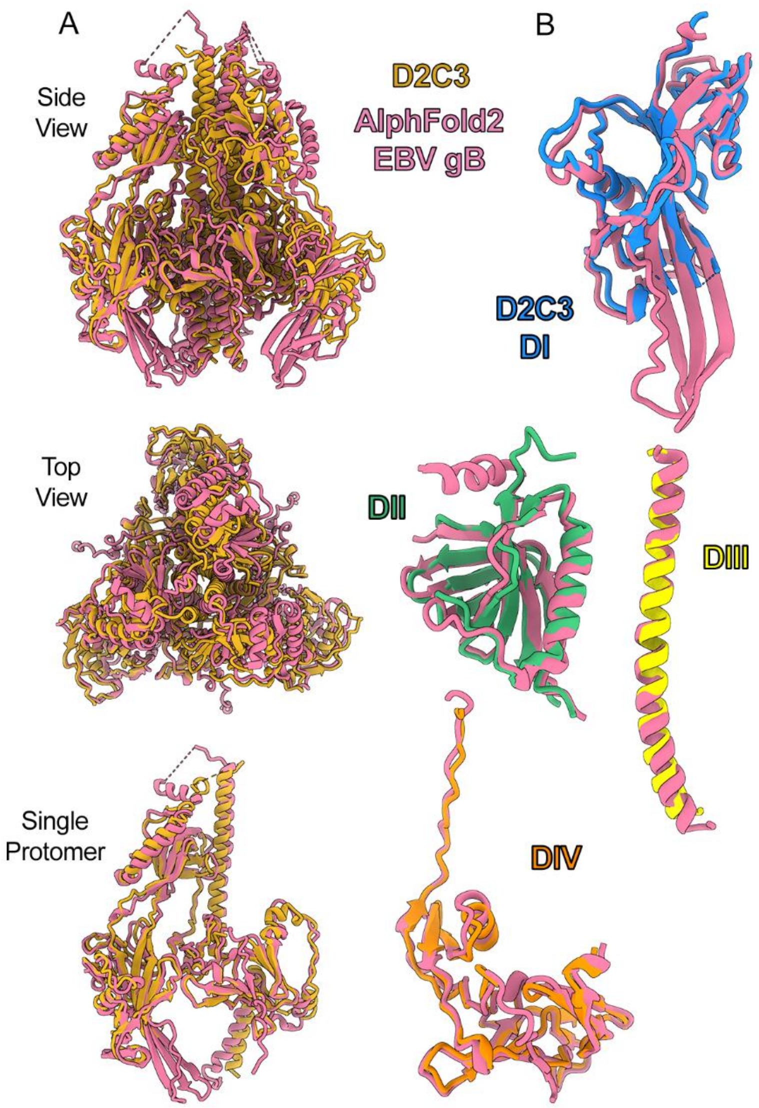


**Supplemental Figure 9 | Alignment of D2C3 structure with the AlphaFold2 model of prefusion EBV gB**. (A) D2C3 (orange) aligned with the EBV gB prefusion model (AlphaFold2, pink). (B) Pairwise alignments of DI–DIV between D2C3 and the EBV gB prefusion model. D2C3 is colored by domain as in Supplementary Fig. S1. The EBV gB prefusion model is colored as in (A). All alignments were performed using the MatchMaker function in ChimeraX[7]. Individual domains were aligned based on residues within the respective domain.

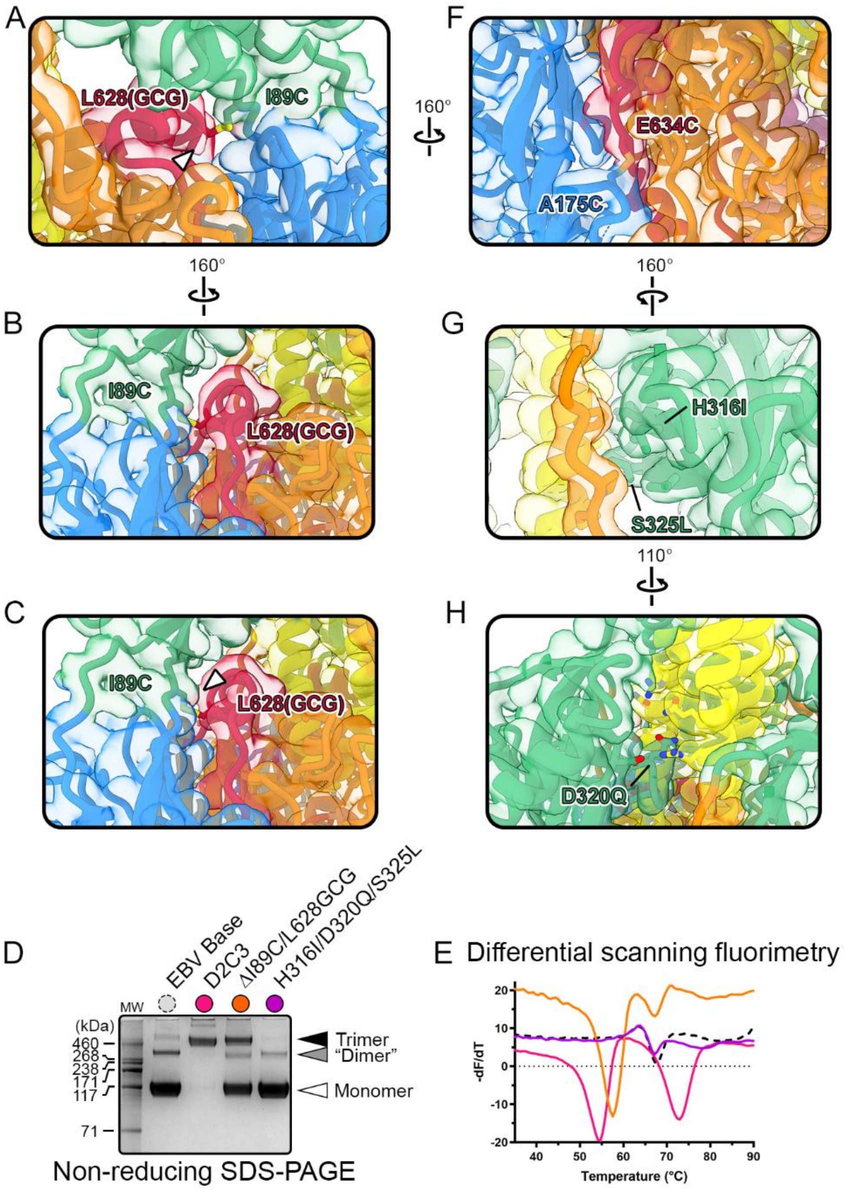


**Supplemental Figure 10 | I89C/L628(GCG) is crucial for prefusion gB stabilization.** (A–B) Zoomed in view of the map-to-model fit of D2C3 near the I89C/L628(GCG) substitutions. (C) Same view as (B), but with a lower cryo-EM map threshold to reveal additional features. The white arrows in (A) and (C) denote an extra map feature observed for the L628(GCG) residues. (D) Non-reducing SDS-PAGE analysis of gB variants. (E) DSF analysis of gB variant thermostability, colored as in (D). (F–H) Zoomed in view of the map-to-model fit of D2C3 near (F) the A175C/E634C substitutions, (G) the H316I and S325L substitutions, and (H) the D320Q substitution. The cryo-EM map is transparent and colored by domain, as in Supplementary Fig. S1. Key residues are shown as sticks. Nitrogen atoms are colored blue, oxygen atoms are colored red, and sulfur atoms are colored yellow.

**Table S4 | Demographics for human sera donors used in depletion studies.**

| Gender | Age | Race | Status |
|---|---|---|---|
| Male | 46 | Black | Normal, Healthy |
| Male | 57 | Black | Normal, Healthy |
| Male | 64 | Black | Normal, Healthy |
| Male | 50 | Black | Normal, Healthy |
| Male | 42 | Black | Normal, Healthy |
| Male | 41 | Black | Normal, Healthy |
| Male | 40 | Black | Normal, Healthy |
| Male | 58 | Black | Normal, Healthy |
| Male | 32 | Black | Normal, Healthy |
| Male | 58 | Black | Normal, Healthy |
| Male | 57 | Black | Normal, Healthy |
| Male | 47 | Black | Normal, Healthy |
| Male | 65 | Caucasian | Normal, Healthy |
